# Genetic Profiling of Protein Burden and Nuclear Export Overload

**DOI:** 10.1101/2020.02.25.962068

**Authors:** Reiko Kintaka, Koji Makanae, Hisaaki Kato, Shinsuke Ohnuki, Yoshikazu Ohya, Brenda Andrews, Charles Boone, Hisao Moriya

**Affiliations:** Donnelly Center for Cellular and Biomolecular Research, Department of Medical Genetics, University of Toronto; Research Core for Interdisciplinary Sciences, Okayama University, Okayama, Japan; Faculty of Agriculture, Okayama University, Okayama, Japan; Graduate School of Frontier Sciences, University of Tokyo, Japan; RIKEN Center for Sustainable Resource Science, Wako, Saitama, Japan; Graduate School of Environmental and Life Science, Okayama University

## Abstract

Overproduction (op) of proteins triggers cellular defects. One of the defined consequences of protein overproduction is the protein burden/cost, which is produced by an overloading of the protein synthesis process. However, the physiology of cells under a protein burden is not well characterized. We performed genetic profiling of protein burden by systematic analysis of genetic interactions between GFP-op, surveying both deletion mutants of nonessential genes and temperature-sensitive mutants of essential genes, in the budding yeast *Saccharomyces cerevisiae*. To dissect interactions specific to the protein burden, we also performed genetic profiling in cells with overproduction of triple-GFP (tGFP), and the nuclear export signal-containing tGFP (NES-tGFP). The mutants specifically interacted with GFP-op were suggestive of unexpected connections between actin-related processes like polarization and the protein burden, which was supported by morphological analysis. The tGFP-op interactions suggested that this protein probe overloads the proteasome, probably through the formation of intracellular aggregates, whereas those that interacted with NES-tGFP involved genes encoding components of the nuclear export process, providing a resource for further analysis of the protein burden and nuclear export overload.

## Introduction

Extreme overproduction of a gratuitous protein that has no cellular function causes growth defects, which, at least in part, appears to be caused by overloading the cellular resources for protein synthesis (Dong *et al.*, 1995; Snoep *et al.*, 1995; Stoebel *et al.*, 2008; Makanae *et al.*, 2013; Shah *et al.*, 2013; Kafri *et al.*, 2016; Moriya, 2015; Eguchi *et al.*, 2018). This phenomenon is called the protein burden/cost and has been extensively studied in the budding yeast *Saccharomyces cerevisiae*, a model eukaryotic cell. Limiting functions defining the protein burden are thought to be the translational process upon nitrogen limitation, and the transcriptional process upon phosphate limitation (Kafri *et al.*, 2016). The protein burden itself initially appears to be a relatively simple phenomenon, but little is known about the physiological conditions and cellular responses triggered by the protein burden.

To trigger the protein burden, a protein must be produced at a level sufficient to overload protein production resources (Moriya, 2015; Eguchi *et al.*, 2018). This can happen only if the protein is otherwise harmless. Fluorescent proteins, such as EGFP, Venus, and mCherry, do not have any physiological activity in yeast cells and thus are considered gratuitous proteins. Therefore, these fluorescent proteins are believed to be produced at the highest possible levels in yeast cells, and their overproduction triggers a protein burden (Makanae *et al.*, 2013; Kafri *et al.*, 2016; Eguchi *et al.*, 2018; Farkas *et al.*, 2018). Modifications to EGFP, such as adding a degradation signal, misfolding mutations, or adding localization signals, reduces its expression limit, probably because these modifications overload limited resources for the degradation, folding, and localization processes, respectively (Geiler-samerotte *et al.*, 2010; Makanae *et al.*, 2013; Kintaka *et al.*, 2016; Eguchi *et al.*, 2018).

A recent study isolated a group of deletion strains in which growth defects upon overproduction of yEVenus are exacerbated (Farkas *et al.*, 2018). Through the analysis of these strains, and conditions exacerbating the protein burden, the authors concluded that Hsp70-associated chaperones contribute to the protein burden by minimizing the damaging impact of the overproduction of a gratuitous protein. Chaperone genes, however, constitute only a relatively small fraction of the deletion strains isolated in the study, and thus the protein burden may impact numerous other processes.

In this study, we surveyed genetic interactions between mutant strains and high levels of GFP overproduction (GFP-op) to genetically profile cells exhibiting this phenomenon. To isolate mutant sets showing positive and negative genetic interactions with the protein burden, we used a condition causing significant growth defects due to high GFP-op from the *TDH3* promoter (TDH3_pro_) on a multi-copy plasmid. In addition to a deletion mutant collection of non-essential genes, we surveyed temperature-sensitive (TS) mutant collections of essential genes. We performed a strict statistical evaluation to isolate mutants showing robust genetic interactions with high confidence.

We also attempted to distinguish between the protein burden and other process overloads by surveying genetic interactions between those mutant strains and a triple-GFP (tGFP) with a nuclear export signal (NES-tGFP). NES-tGFP triggers growth defects at a lower expression level than unmodified tGFP (Kintaka *et al.*, 2016). If the protein burden can only be triggered by a harmless protein like GFP, mutants harboring genetic interactions with tGFP-op should be different from those with NES-tGFP-op, and the comparison of those mutants will identify consequences specific to the protein burden. Moreover, mutants harboring negative genetic interactions should contain limiting factors of the nuclear export and essential factors affected by the overloading of nuclear export.

## Results

### Isolation of mutants that have genetic interactions with GFP-op

To isolate mutants genetically interacting with GFP-op, we performed a synthetic genetic array (SGA) analysis (Baryshnikova *et al.*, 2010) **(Figure 1A**). As a query strain, we overproduced GFP (yEGFP) (Cormack and Bertram, 1997) under the control of TDH3_pro_ on the multi-copy plasmid pTOW40836 (**Figure 1B**). This plasmid contains two selection markers (*URA3* and *leu2-89*), and the copy number can be controlled by the culture conditions. The copy numbers of this plasmid under –Ura and –Leu/Ura conditions are around 8 and 30 copies per cell, respectively (Eguchi *et al.*, 2018). While a strain harboring this plasmid shows growth defects even under –Ura conditions (**Figure 1C**), the strain shows more growth defects under –Leu/Ura conditions (**Figure 1C**), presumably because the copy number increase leads to an increase in GFP production, and probably causes a higher protein burden (Eguchi *et al.*, 2018).

**Figure 1.**
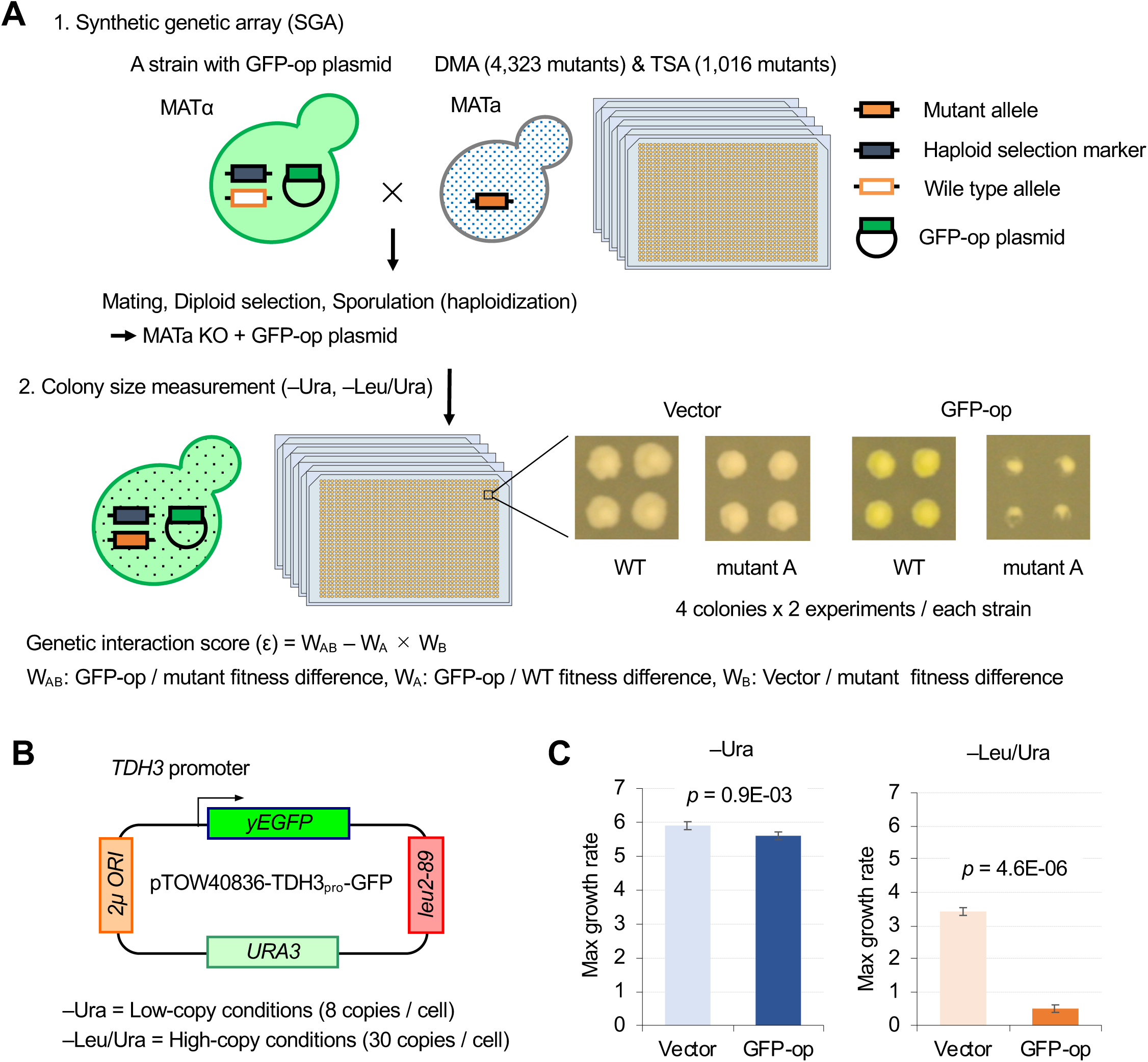
Experimental scheme of genetic interaction (GI) analysis. **A.** Each mutant from a deletion mutant array (DMA) and a temperature-sensitive mutant array (TSA) was combined with GFP overproduction (GFP-op) using the synthetic genetic array (SGA) method (Baryshnikova *et al.*, 2010). The colony size of each derivative strain grown on synthetic complete (SC)–Ura and SC–Leu/Ura plates was measured to calculate a genetic interaction (GI) score (ε). Four colonies were analyzed for each strain, and the entire experiment was duplicated. **B.** The structure of the plasmid used to overexpress GFP. The plasmid copy number, and thus the expression level of GFP, can be changed by changing the growth conditions. **C.** Effect of GFP production on growth under each condition. The maximum growth rate was measured in liquid culture. The average, standard deviation (error bar), and *p*-value of Student’s t-test for four independent experiments are shown.

We examined an array of 4,323 deletion mutants in nonessential genes (DMA) and an array of 1,016 conditional temperature-sensitive mutants (TSA) (Costanzo *et al.*, 2016). For each mutant strain, we calculated genetic interaction (GI) scores (ε) from the analysis of four colonies under both –Ura and –Leu/Ura conditions, in duplicate (**Data S1**). After thresholding by the variation in colony size (*p* < 0.05), we compared GI scores between duplicates (**Figure 2A, Figure 2-S1**). The reproducibility of the DMA experiments was lower in –Ura conditions (*r* = 0.17), whereas it was higher in –Leu/Ura conditions (*r* = 0.36). The reproducibility of the TSA experiments was higher in both – Ura and –Leu/Ura conditions (*r* = 0.42 and 0.53). Thus, the conditions which cause severe growth defects produce the most reproducible GI scores.

**Figure 2.**
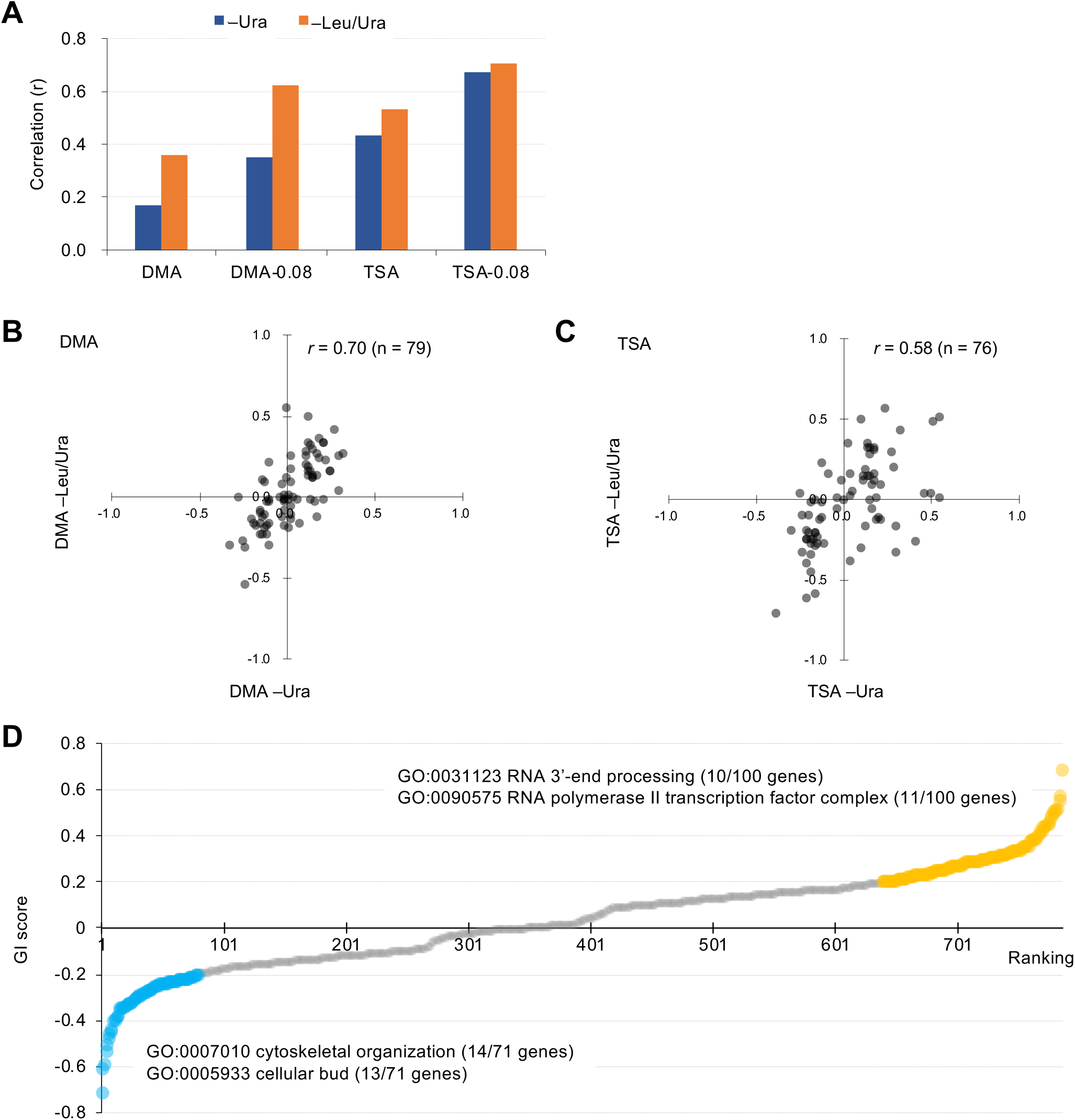
Characteristics of GI scores. **A**. Pearson correlation coefficient (*r*) of GI scores from experimental duplicates. DMA and TSA: comparison of all GI scores of duplicates obtained by the GI analysis using DMA and TSA. DMA-0.08 and TSA-0.08: comparison of GI scores of duplicates with value > |0.08| obtained by the GI analysis using DNA and TSA. **Figure 2S-S1** shows an independent comparison. **B** and **C**. Comparison of average GI scores of DMA (**B**) and TSA (**C**) mutants both with GI scores in the duplicates > |0.08| under –Ura and –Leu/Ura conditions. **C**. GI score (ε) of mutants isolated ordered by score ranking. Mutants with low (<0.2) and high (>0.2) scores are shown in light blue and orange, with enriched GOs in those mutants. The score in –LU is shown.

To more confidently identify mutants showing strong GIs, we set a threshold in each replicate (ε > |0.08|). Using this threshold increased reproducibility, especially in the DMA experiments (*r* = 0.35 in –Ura, *r* = 0.62 in –Leu/Ura, **Figure 2A**). We first selected mutants with ε > |0.08| in each replicate and then calculated their average GI scores between the duplicates as **summarized in Figure 2-S2, 2-S3**. Because GI scores between –Ura (low-level GFP-op) and –Leu/Ura (high-level GFP-op) conditions were highly correlated (*r* = 0.70 and 0.58, **Figure 2B and 2C**), this procedure identified high-confidence mutants with GIs with GFP-op.

Farkas *et al.* surveyed GIs between deletion mutants and the overproduction of yEVenus. The GI scores obtained by our analysis did not show correlation with those from the Farkas study (*r* = –0.01 and –0.07, **Figure 2-S3A, B**). This may be because of the weak reproducibility observed in lower overproduction conditions (**Figure 2S-3C, D**). Moreover this overlap analysis only involved nonessential genes and the Farkas study used a relatively weaker *HSC82* promoter (HSC82_pro_), in medium comparable to our –Ura condition, in which the GFP-op from HSC82_pro_ on pTOW40836 caused a very minor growth defect in –Ura conditions (**Figure 2S-3E, F**). Indeed, our conditions produced more variance in the GI scores and thus identified more mutants showing stronger GIs (**Figure 2-S3A, B)**, and we found that negative GIs of 6 out of 7 deletion mutants from our screening were confirmed by independent growth measurements in the liquid medium, while all six mutants isolated by the previous study (Farkas *et al.*, 2018) were not (**Figure 2-S4A, B**).

During the screening, we noticed that a group of TS mutants showed greater growth defects under –Leu/Ura conditions than under –Ura conditions in the vector control experiments (**Figure 2-S5**). The gene ontology (GO) term “DNA replication preinitiation complex [GO:0031261]” was significantly over-represented in the mutated genes (seven genes, *p* = 1.47E–05). **Figure 2-S5A** shows the normalized colony size differences of the 18 mutants analyzed in the TSA corresponding to the genes categorized in GO:0031261. 6 out of 18 mutants showed more than 2U decrease in their colony sizes, whereas the average of all TS mutants showed 0.002U (Rep1) and 0.003U (Rep2). The vector copy number is more than 100 copies per cell under –Leu/Ura conditions (Makanae *et al.*, 2013; Eguchi *et al.*, 2018). This high copy number probably produces limitations of the replication initiation complex by sequestering the complex to the replication origins of the plasmids. Some negative factors on the plasmid, like TDH3_pro_-GFP, restrict the plasmid copy number due to a genetic tug-of-war effect (Moriya *et al.*, 2006), and the plasmid thus may not trigger the limitation of the replication initiation complex. This situation may lead to a bias toward the isolation of mutants in the replication initiation complex with positive GIs with plasmids containing toxic elements, especially under –Leu/Ura conditions.

### Mutations aggravating or mitigating GFP-op triggered growth defects

To understand which processes are affected by GFP-op, we performed enrichment analysis targeted toward isolating mutants with stronger GIs (ε > |0.2|) under –Leu/Ura conditions, as the results obtained under these conditions were more reproducible (**Figure 2A, Data S2**). We designated the negatively interacting genes and mutants “GFP-op_negative” and the positively-interacting genes and mutants “GFP-op_positive.” The GFP-op_negaitive 71 genes (79 mutants) were significantly enriched in GO categories related to cytoskeletal organization and polarization (**Figure 2D, Table S1**). **Figure 3A** shows the GI scores under –Leu/Ura conditions of all 45 alleles of the GFP-op_negative genes categorized in GO as “cellular bud [GO:0005933].” Most of the mutants showed negative GIs, and 16 out of 45 showed average scores of less than – 0.2.

**Figure 3.**
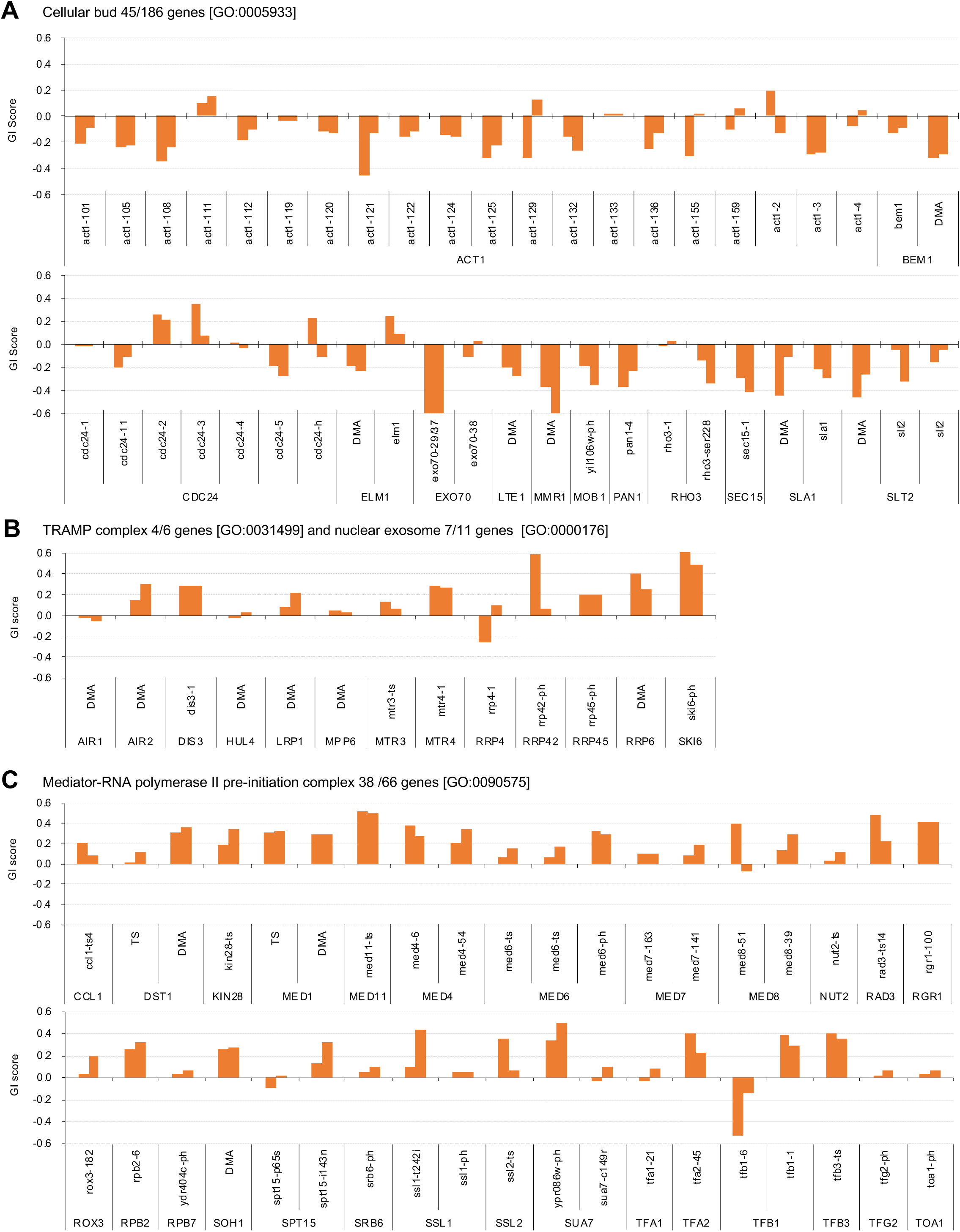
Independent GI scores (ε) of genes enriched in GO categories in GFP_negative and GFP_positive genes. **A.** GI scores of mutants isolated as GFP_negative genes annotated with the GO term “cellular bud [GO:0005933].” **B.** GI scores of mutants annotated with the GO terms “TRAMP complex [GO:0031499]” and “nuclear exosome [GO:0000176].” **C.** GI scores of mutants annotated with the GO term “Mediator-RNA polymerase II preinitiation complex [GO:0090575].” GI scores under –Leu/Ura conditions are shown.

One hundred GFP-op_positive genes (100 mutants) were enriched in genes involved in RNA 3′-end processing and the transcription factor complex (**Figure 2D, Table S1**). Among the factors in the RNA 3′-end processing, the subunits in the “TRAMP complex [GO:0031499]” and “nuclear exosome [GO:0000176]” were isolated as GFP-op_positive genes. **Figure 3B** shows the GI scores under –Leu/Ura conditions of the mutants of the TRAMP complex and the nuclear exosome subunits. 7 out of 13 mutants showed positive GIs with average scores greater than 0.2. Among the transcription factor complex, subunits of the “mediator-RNA polymerase II preinitiation complex [**GO:0090575**]” were specifically isolated. **Figure 3C** shows the GI scores under –Leu/Ura conditions of the mutants of the mediator-RNA polymerase II preinitiation complex subunits. In total, 20 out of 38 mutants showed positive GIs with average scores greater than 0.2.

### Investigation of GFP expression levels of mutants

We next investigated GFP expression levels of the GFP-op_negative and GFP-op_positive mutants. To obtain the GFP expression level of each mutant, we measured normalized GFP fluorescence (GFPunit) from the fluorescence intensity of each colony (**Figure 4A**). Of the mutants, 1447 (29%) showed lower GFPunits and 3572 (71%) mutants show higher GFPunits than the average of all mutants (**Figure 4B and 4C, Data S2**). We designated these mutants GFP_H and GFP_L, respectively (**Figure 4B and 4C**).

**Figure 4.**
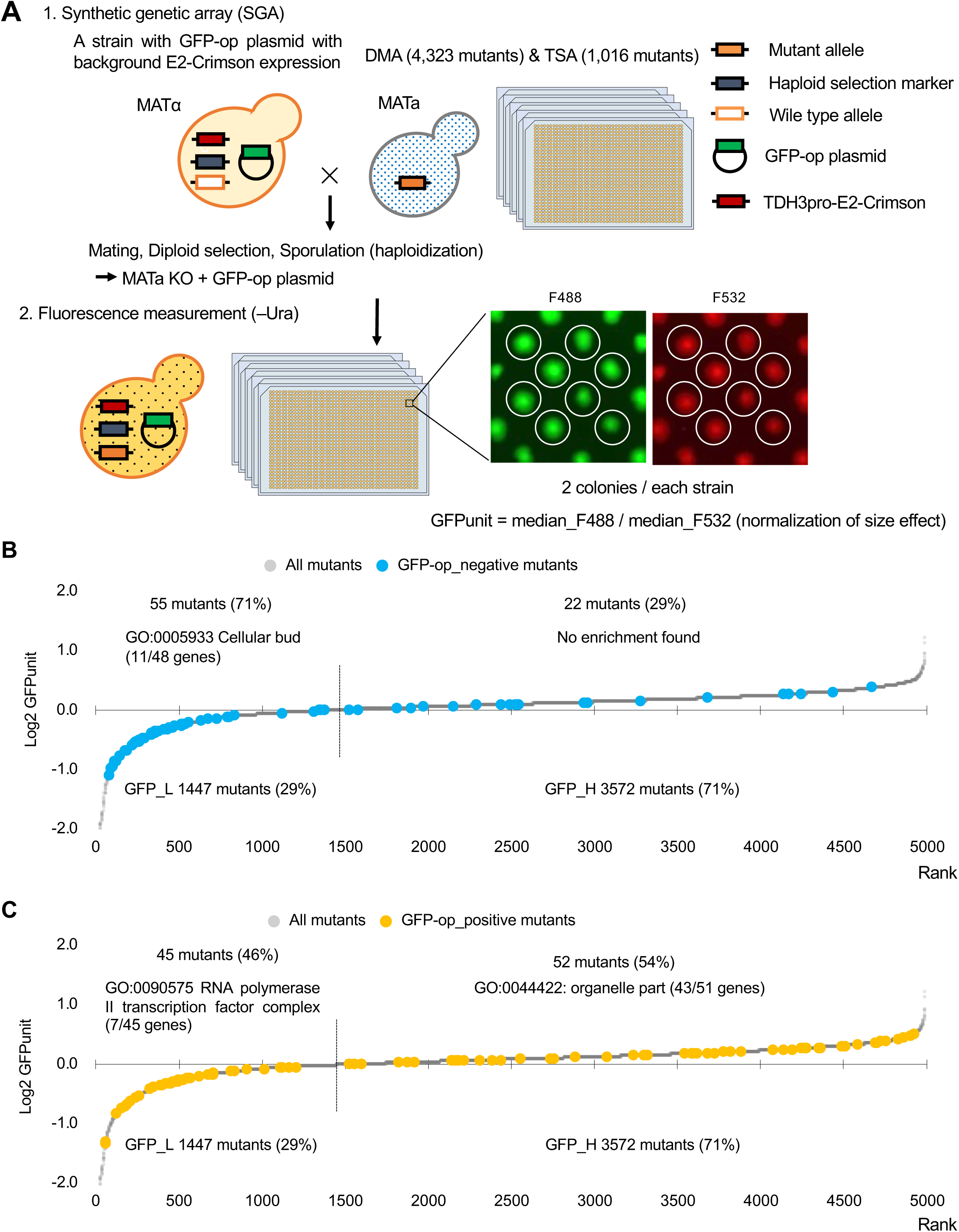
Experimental scheme of GFP expression measurements of mutants. **A.** Each mutant from a deletion mutant array (DMA) and a temperature-sensitive mutant array (TSA) was combined with GFP overproduction (GFP-op) with background E2-Crimson expression, using a synthetic genetic array (SGA) method. The median GFP fluorescence (F488) and E2-Crimson fluorescence (F532) of each colony were measured, and the GFP expression level (GFPunit) of each mutant was calculated by dividing F488 by F532 to normalize colony size. **B.** GFPunits of GFP-op_negative mutants. Mutants with lower and higher GFPunits than the average are designated as GFP_L and GFP_H mutants, respectively. Representative GO terms enriched in GFP_L mutants in GFP-op_negative mutants are shown. **C.** GFPunits of GFP-op_positive mutants. Representative GO terms enriched in GFP_L mutants and GFP_H mutants in GFP-op_positive mutants are shown.

GFPunit can be used to interpret the mechanisms underlying GFP-op_negative and GFP-op_positive mutations as follows: 1) if GFPunit is lower in a GFP_negative mutant, the mutant is considered to be more sensitive to GFP overproduction; 2) if GFPunit is higher in a GFP_negative mutant, the mutant triggers greater GFP overproduction, which may cause more protein burden and growth defects; 3) if GFPunit is lower in a GFP_positive mutant, the mutant triggers lower GFP overproduction, which may cause less protein burden and growth defects; and 4) if GFPunit is higher in a GFP_positive mutant, the mutant is considered to be less sensitive to GFP overproduction.

GFP-op_negative mutants were significantly enriched in GFPunit_L mutants (**Figure 4B**, *p* = 4.7E-11, Student’s t-test). Because 11 out of 13 GFP-op_negative mutants categorized as “cellular bud [GO:005933]” were also GFP_L (**Figure 4-S1A, Table S2**), these mutants seemed to be sensitive to the protein burden. In contrast, GFP-op_positive mutants were only slightly enriched in GFP_L mutants (**Figure 4C**, *p* = 0.013, Student’s t-test). Trends in the distributions of mutants in “TRAMP complex [GO:0031499],” “nuclear exosome [GO:0000176],” and “mediator-RNA polymerase II preinitiation complex [PMID27610567]” were not obvious (**Figure 4-S1B, Table S2**). However, GFP-op_positive and GFP_L mutants were significantly enriched in “RNA polymerase II transcriptional factor complex [GO:0090575],” suggesting that these mutants may simply cause the reduction of GFP production, but not decrease the sensitivity to the protein burden.

### Overproduction of tGFP and NES-tGFP results in GIs with distinct sets of genes

We next analyzed mutants genetically interacting with a GFP containing a nuclear export signal (NES). For this purpose, we used a *PYK1* promoter (PYK1_pro_)-driven tGFP with an NES from PKI **(Figure 5A, Data S3**) because its exclusion from the nucleus has been confirmed (**Kintaka 2016**), and the degree of growth inhibition is similar to that of TDH3_pro_-GFP (**Figure 5B**). We also used PYK1_pro_-tGFP as a control for NES-tGFP (**Figure 5A, Data S4**). Using the same procedures as in the analysis of GFP described above except upper and lower threshold of ε 0.16 and –0.12, we isolated total 714 mutants (695 genes) harboring GIs with either GFP-op, tGFP-op or NES-tGFP-op under –Leu/Ura conditions (**Data S5**). To extract genes that had specific GIs with each condition, we performed clustering analysis using them, which were isolated in at least one of GFP-op, tGFP-op, and NES-tGFP experiments (**Figure 5C, Data S6)**.

**Figure 5.**
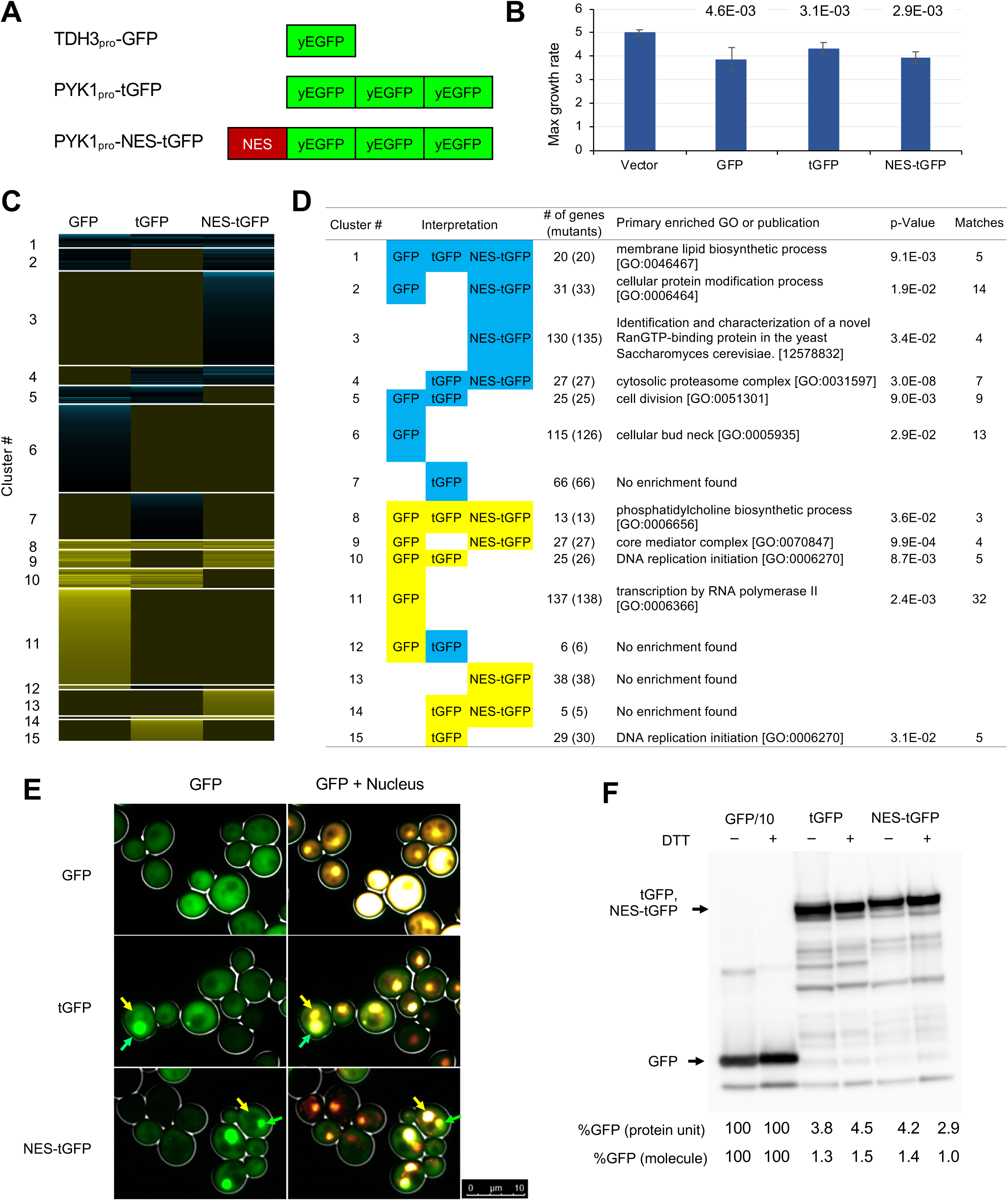
GFP-op harbor GIs with distinct sets of genes from those with tGFP-op and NES-tGFP-op. **A.** Structures and promoters used to overexpress GFP, tGFP, and NES-tGFP. Nucleotide sequences of the three GFPs in tGFP (and NES-tGFP) are different, other to avoid recombination. **B. Maximum growth rates of cells harboring overproduction plasmids** The average, standard deviation, and *p*-value of Student’s t-test for the growth rates of cells with the vector and overproduction plasmids from four independent experiments are shown. Cells were grown in synthetic complete (SC)–Ura medium. **C and D.** Clustering analysis of the mutants having GIs with GFP-op, tGFP-op, and NES-tGFP-op (C), and its characterization (D). **E.** Microscope images of cells overexpressing GFP, tGFP, and NES-tGFP The nucleus was observed using Hoechst 33342 staining. Representative cells with intracellular condensates are indicated by green arrowheads (condensates with GFP fluorescence) and yellow arrowheads (nucleus). **F. Quantification of expression limits of GFP, tGFP, and NES-tGFP** Western blot analysis of total protein from GFP-op (1/10 diluted), tGFP-op, and NES-tGFP cells cultured in SC–Ura medium. Relative GFP levels (protein units) were calculated by measuring the intensities of bands corresponding to the molecular weight of each protein (arrowheads). Note that molar concentration GFP should be divided by three in the case of tGFP and NES-tGFP because they have three times more epitopes for the antibody than GFP.

**Figure 5D** shows the representative GO term or publication for each cluster (the whole data is shown in **Table S3**). Mutants negatively interacting only with NES-tGFP-op (Cluster 3) contained mutants of genes playing a central role in the nuclear protein export (Crm1, Gsp1, Rna1, and Yrb1). GI scores of these mutants were significantly lower in the NES-tGFP-op experiment than in the other two experiments (**Figure 5-S1A**), suggesting that NES-tGFP-op specifically causes growth defects through overloading these limited factors.

Only 12% (81/688) of mutants showed shared GIs between GFP and tGFP. Mutants negatively interacting with tGFP-op and NES-tGFP-op (Cluster 4) were strongly enriched in annotations of “cytosolic proteasome complex [GO:0031597]” (**Figure 5D**). GI scores of mutants in “proteasome complex [GO:0000502]” were significantly lower in the tGFP-op and NES-tGFP-op experiments than in the GFP-op experiment (**Figure 5-S1B**). These results suggest that GFP and tGFP have different characteristics, and tGFP-op triggers proteasome stress. Interestingly, we observed aggregative structures in the cytoplasm of tGFP-op and NES-tGFP-op cells but not in GFP-op cells (**Figure 5E**).

Mutants interacting only with GFP-op (Cluster 6 and Cluster 11) were enriched in genes annotated to “cellular bud neck [GO: 0005935]” and “transcription by RNA polymerase II [GO:0006366],” and their GI scores were significantly lower and higher in tGFP-op experiments than in the other two experiments (**Figure 5-S1C, D**). This observation suggests that these two processes are specifically interacting with the protein burden, and can be only triggered by proteins with very high expression. Expression levels of tGFP and NES-tGFP which caused growth defects were less than 2% of that of GFP (**Figure 5F**).

### GFP-op affects actin distribution

The above results indicate that GFP-op, i.e. the protein burden, could affect actin functions. We thus performed a morphological analysis of cells under GFP-op with a high-throughput image-processing system (CalMorph) (Ohtani *et al.*, 2004). We used non-fluorescent GFP mutant (GFPy66g) for this analysis because strong GFP fluorescence affects the observation of the cell shape with FITC-ConA. We also analyzed the cells overexpressing Gpm1 and a catalytically negative Gpm1 mutant (Gpm1-m) whose overexpression is considered to cause the protein burden (Eguchi *et al.*, 2018). Cells were cultured under SC–Ura conditions. Among obtained 502 morphological parameters, only four parameters showed significant differences over the vector control, and three of them (A120_A1B, ACV7-1_A, and A122_A1B) were actin-related parameters (**Figure 6A-D**). **Figure 6E** shows the interpretation of the morphology of GFP-op cells. The cells contained increased actin patch regions, supporting the idea that the protein burden interacts with actin function.

**Figure 6.**
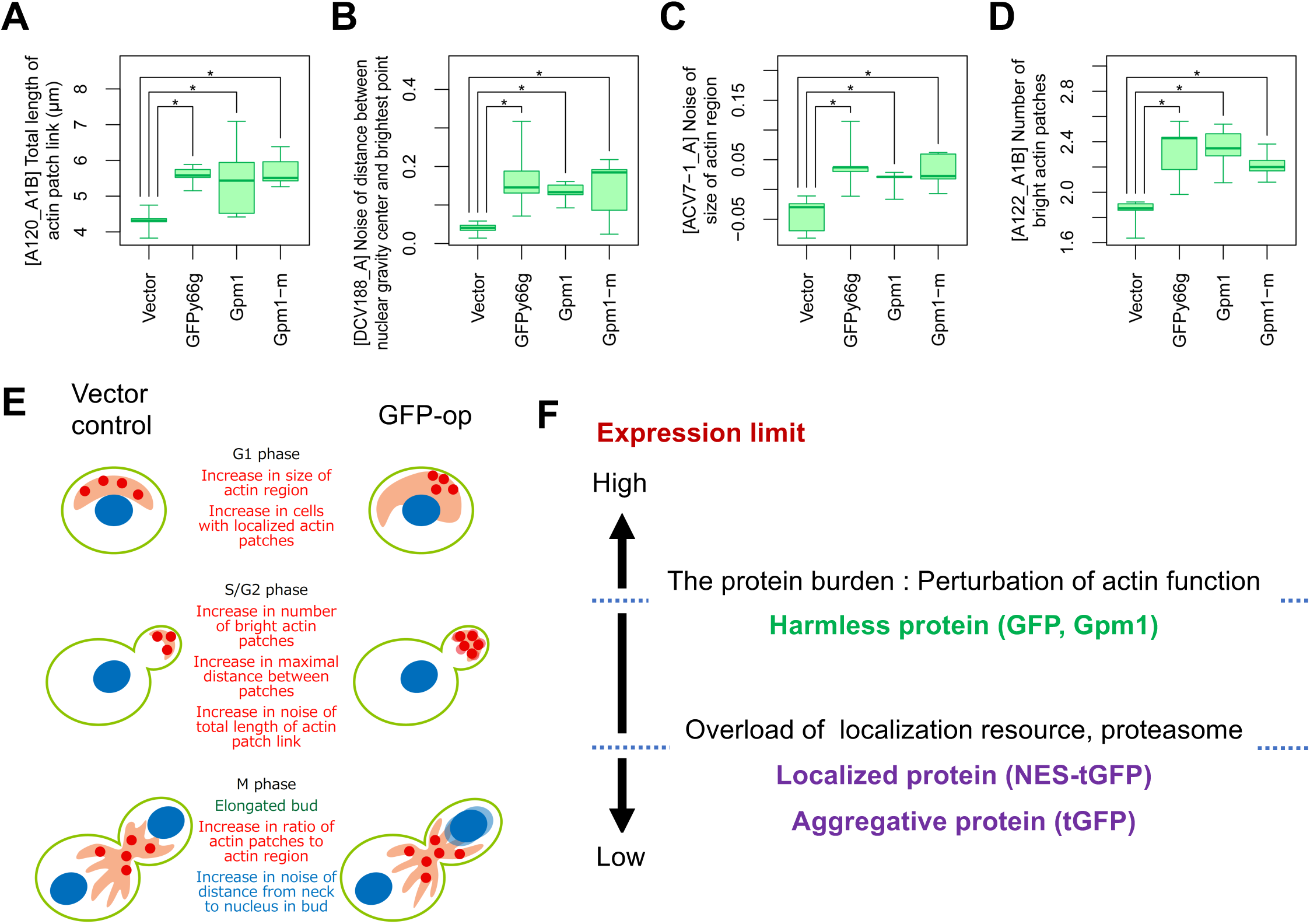
Morphological analysis of the cells overexpressing gratuitous proteins. **A-D**. Morphological parameters significantly different all in the cells overexpressing GFPy66g, Gmp1, and Gpm1-m cells over the cells with the vector control. *: FDR = 0.01 by Wald test. To overexpress GFPy66g, Gpm1, and Gpm1-m, pTOW40836-TDH3_pro_-GFPy66g, pTOW40836-TDH3_pro_-Gpm1, and pTOW40836-TDH3_pro_-Gpm1-m were used. **E.** Interpretation of the morphology of GFP-op cells according to the morphological parameters significantly different from the vector control. **F.** Dissection of the consequence of protein overexpression by the expression limits. Only otherwise harmless protein could cause the protein burden, which is associated with the perturbation of actin function.

## Discussion

In this study, we genetically profiled the consequences of protein overproduction using GFP as a model gratuitous protein and NES-tGFP as a transported model protein. We confirmed our prediction that the overproduction of NES-containing protein (NES-tGFP) overloads the amount of limiting nuclear-export factors (Kintaka *et al.*, 2016). Overproduction of NES-tGFP had strong negative GIs with mutants in the major nuclear export factors (Crm1, Gsp1, Rna1, and Yrb1; **Figure 5D and Figure 5-S1A**). tGFP-op (and NES-tGFP-op) had negative GIs with mutants in proteasome components but GFP-op did not (Figure 5D and Figure5-S1B). We observed aggregation of tGFP and NES-tGFP (Figure 5E). Therefore, an overproduction of a protein that contains tandem repeats might induce aggregations, which are degraded by the proteasome.

A comparison of mutants interacting with overproduction of three model proteins led to the isolation of mutants which specifically interact with GFP-op (Figure 5). The three model proteins caused growth defects with different expression levels (Figure 5B and 5F). The GFP level is considerably higher than the levels of tGFP and NES-tGFP, and its expression is the highest of all proteins in yeast (Eguchi *et al.*, 2018), suggesting that overproduction of GFP causes growth defects because of the protein burden. As the protein burden should be triggered by the overproduction of otherwise non-harmful proteins like GFP (Moriya, 2015), these mutants should either exacerbate or mitigate the protein burden. The protein burden is considered to be growth defects occurring as a result of the overloading of protein synthesis processes (Kafri *et al.*, 2016). In contrast to the expectation that mutants in those processes exacerbate the protein burden, the mutants isolated did not show any GO term enrichment in those processes but showed enrichment in actin-related processes like “cytoskeletal organization” or “cellular bud” (Figure 2D). Morphological analysis of cells also supported that GFP-op affects normal actin functions (Figure 6). This relationship might be a result of the long-known connection between actin and translational machinery (Kim and Coulombe, 2010); the protein burden-triggered growth defects might involve the perturbation of the actin cytoskeleton via translational factors like eEF1A which can bundle actin fibers (Munshi *et al.*, 2001). Mutations that mitigate the protein burden indeed enriched genes involved in protein synthesis, especially the transcriptional processes “RNA 3′-end processing” and “RNA polymerase II transcription factor complex” (Figure 2D). Because GFP expression levels in those mutants were lower than average (Figure 4C), those mutants might simply reduce the transcription of the GFP transcript itself.

It is thought that only harmless proteins can be produced up to “the ultimate expression level” to cause the protein burden because harmful proteins should cause cellular defects at lower expression levels (Moriya, 2015). Those defects should be triggered by overloading more limited cellular resources, such as those used for folding and transport, accelerated non-specific interactions, or untimely activation of pathways (Moriya, 2015). Our study here supported this idea through the following observations: 1) tGFP (and NES-tGFP) consists of aggregates in the cell and thus could cause proteostasis stress (Figure 5C); 2) NES-tGFP further uses the protein export machinery; 3) genetic profiling suggested that tGFP-op and NES-tGFP-op overload the proteasome and protein export machinery (**Figure 5D**); 4) expression levels of tGFP and NES-tGFP which cause growth defects are far lower than that of GFP (Figure 5B and 5F); and 6) GFP-op isolated specific mutants that were not isolated in tGFP-op and NES-tGFP. Figure 6F provides a schematic model summarizing this idea. Only harmless proteins like GFP can be produced up to the ultimate expression levels that cause the protein burden, which seems to be related to actin functions. Other proteins, localized or aggregative, can be produced at far lower levels than the level which causes the protein burden because their overproduction overloads localization or protein degradation resources which are more limited than the protein synthesis resource.

In conclusion, our genetic profiling successfully investigated the consequences of overproduction: overload of protein synthesis, nuclear export, and the proteasome. Mutants isolated in this study will be useful resources for further investigations into the general consequences of protein overproduction, especially the overloading of cellular processes.

## Legends

**Figure 2-S1.**
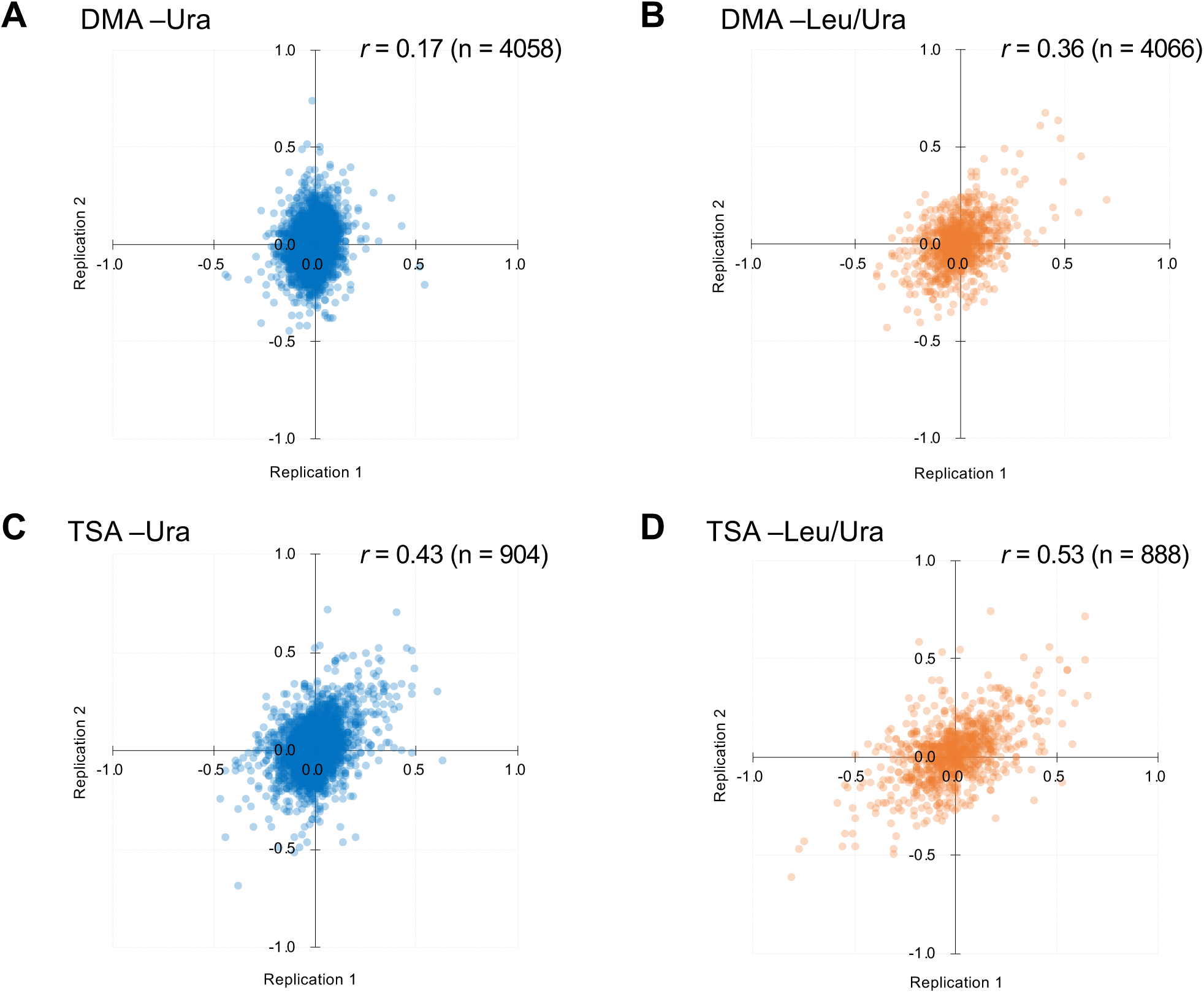
Comparison of GI scores from experimental duplicates. **A-C** indicates the conditions and mutants used.

**Figure 2-S2.**
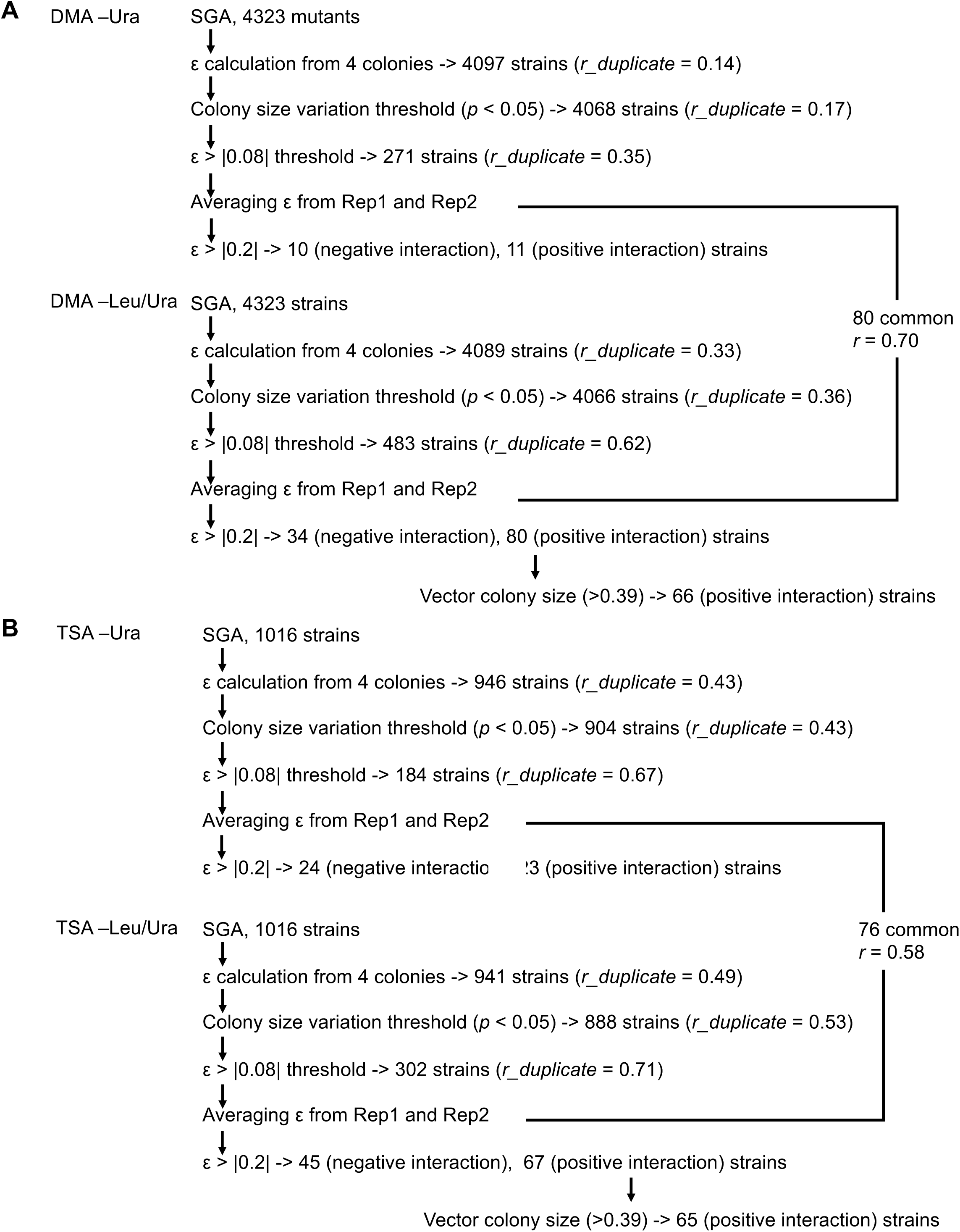
Scheme to isolate mutants showing GIs of high confidence with GFP-op. **A**: analysis with DMA, and **B**: analysis with TSA.

**Figure 2-S3.**
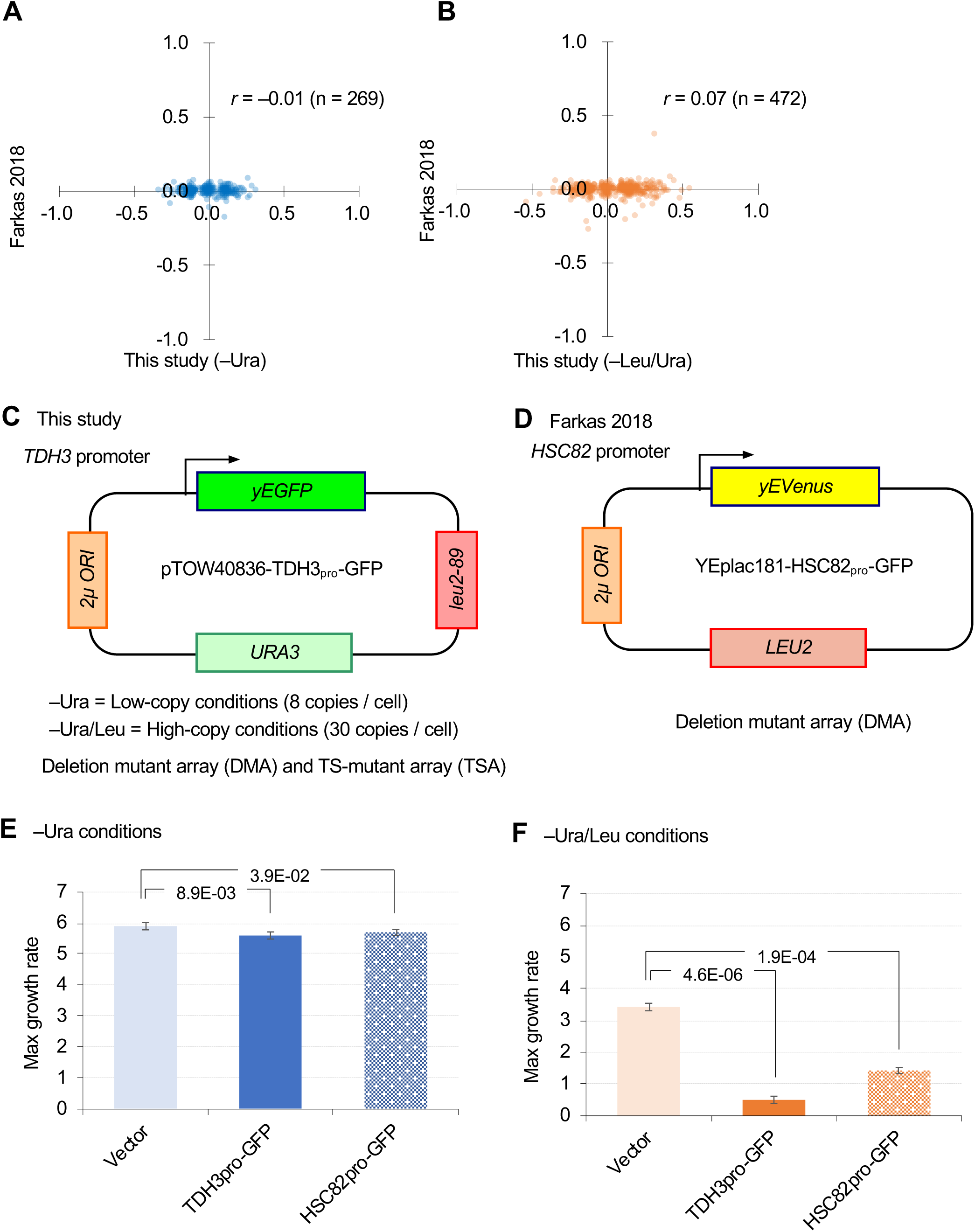
Comparison of GI analyses in this study and a previous study (Farkas *et al.*, 2018). **A** and **B**. Comparison of GI scores of DMA mutants isolated by ε > |0.08| thresholding in this study. **C** and **D**. Plasmids and conditions used in this study and the previous study. **E** and **F**. Effects of overproduction of GFP from TDH3 promoter (TDH3_pro_) and HSC82 promoter (HSC82_pro_) on the plasmid pTOW40836 in –Ura (C) and –Leu/Ura (D) conditions. The average, standard deviation (error bar), and *p*-value of Student’s t-test are shown.

**Figure 2-S4.**
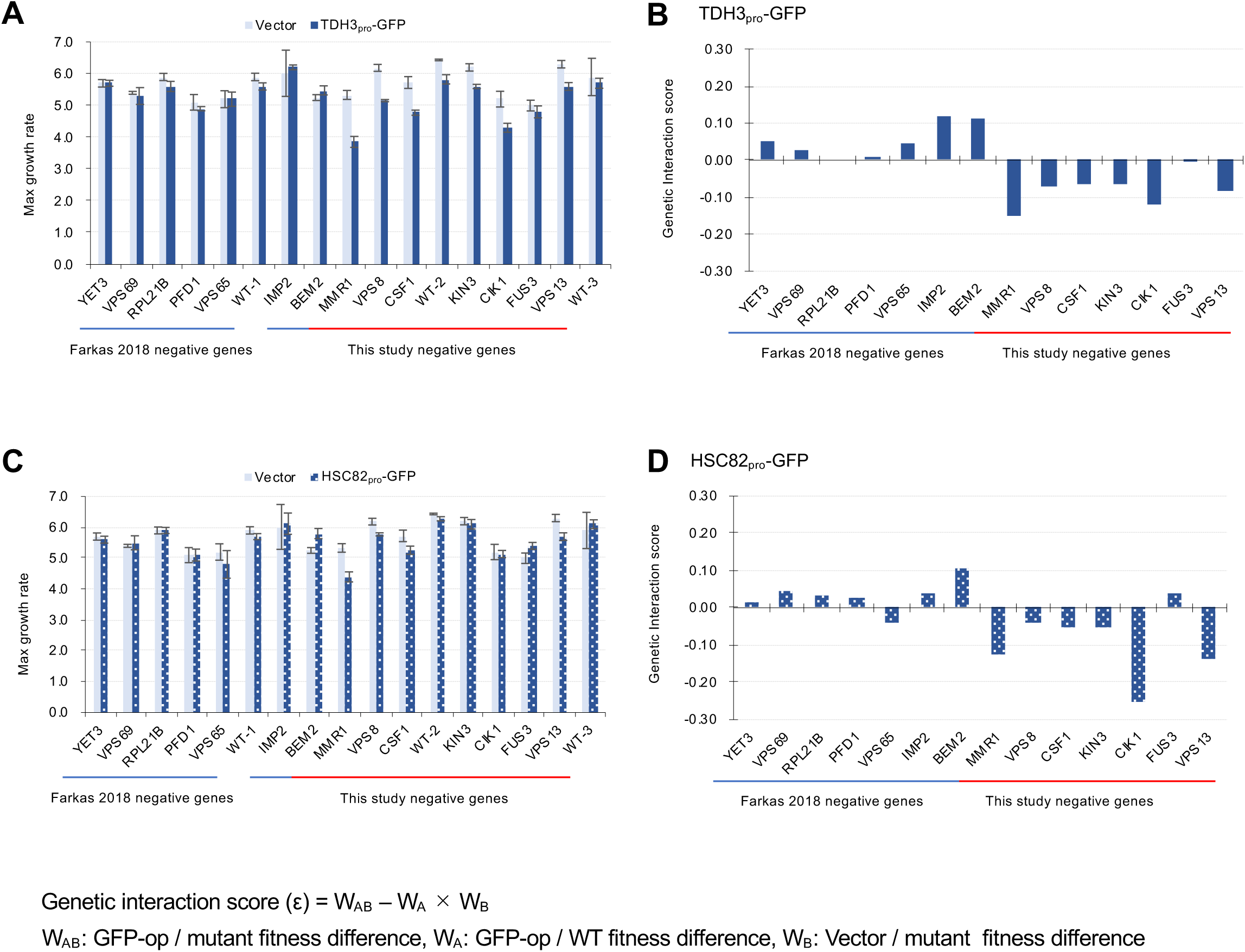
Verification of GIs with independent liquid growth measurement. **A** and **C**. Maximum growth rates of mutant cells with TDP3_pro_-GFP (A) and HSC82 pro-GFP (C) plasmids in synthetic complete (SC)–Ura medium. Average and standard deviation (error bar) of four independent experiments is shown. **B** and **D**. GI scores of mutants and GFP-op from TDH3 _pro_ (**B**) and HSC82 _pro_ (**D**). GI score was calculated as follows: GI score (ε) = W_AB_ – W_A_ × W_B._ Where W_AB_: Gm/Vw, W_A_: Gw/Vw, W_B_: Vm/Vw Gw: Average max growth rate of GFP-op_wild type (four independent measurements) Gm: Average max growth rate of GP-op_mutant (four independent measurements) Vw: Average max growth rate of Vector_wild type (four independent measurements) Vm: Average max growth rate of Vector_mutant (four independent measurements)

**Figure 2-S5.**
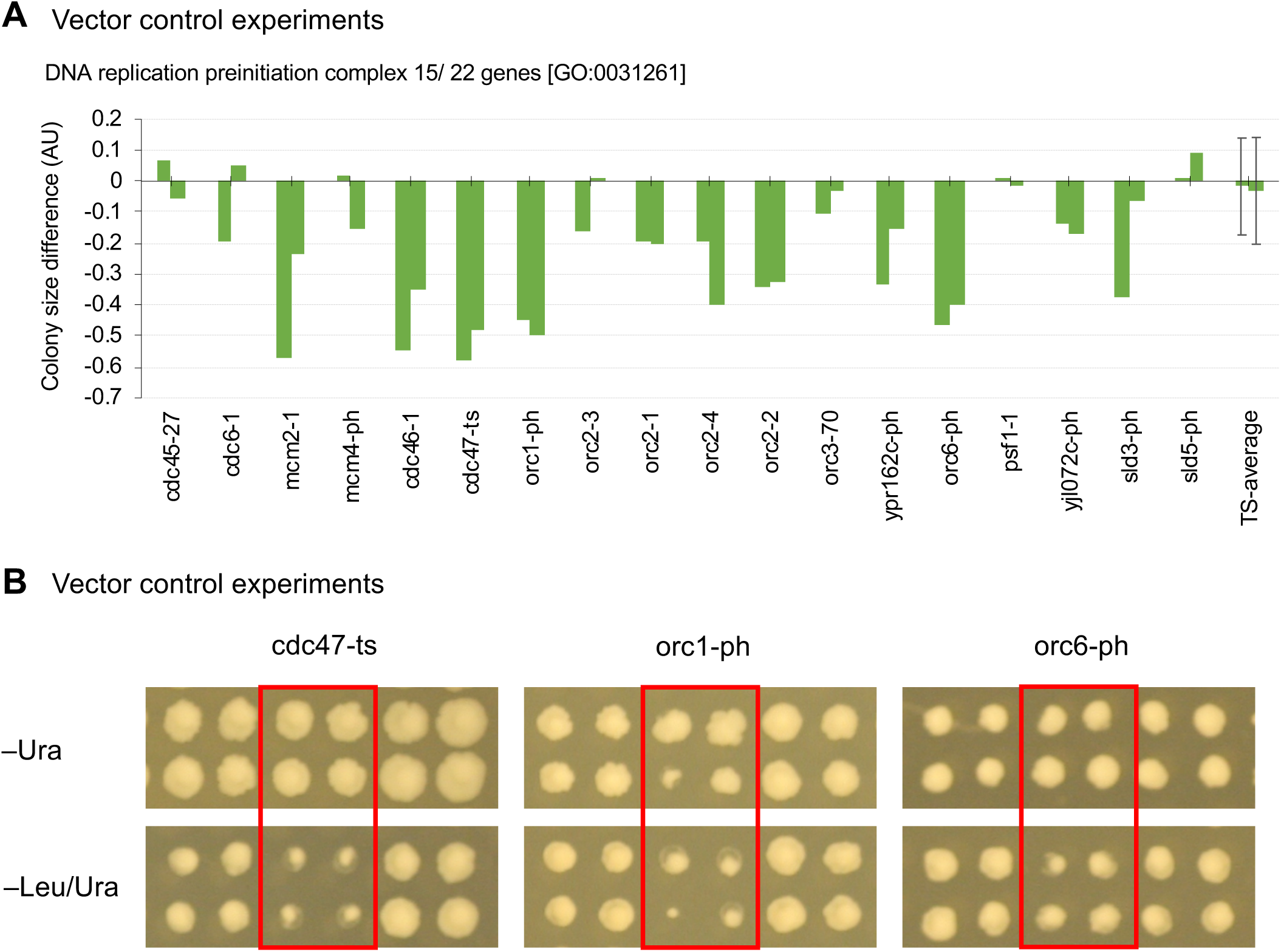
Mutants of replication initiation complex specifically show growth defects in the high-copy conditions. **A.** Colony size differences of vector control experiment of genes categorized as GO categories “DNA replication preinitiation complex [GO:0031261]” on synthetic complete (SC)–Ura and SC–Leu/Ura plates. AU = colony size on –Leu/Ura plate/colony size on –Ura plate. AUs of each mutant from duplicated experiments are shown. The average AU and standard deviation (error bar) of TS mutants are shown. **B.** Representative mutants showing growth defects under high-copy conditions (– Leu/Ura) in the vector control experiments.

**Figure 4-S1.**
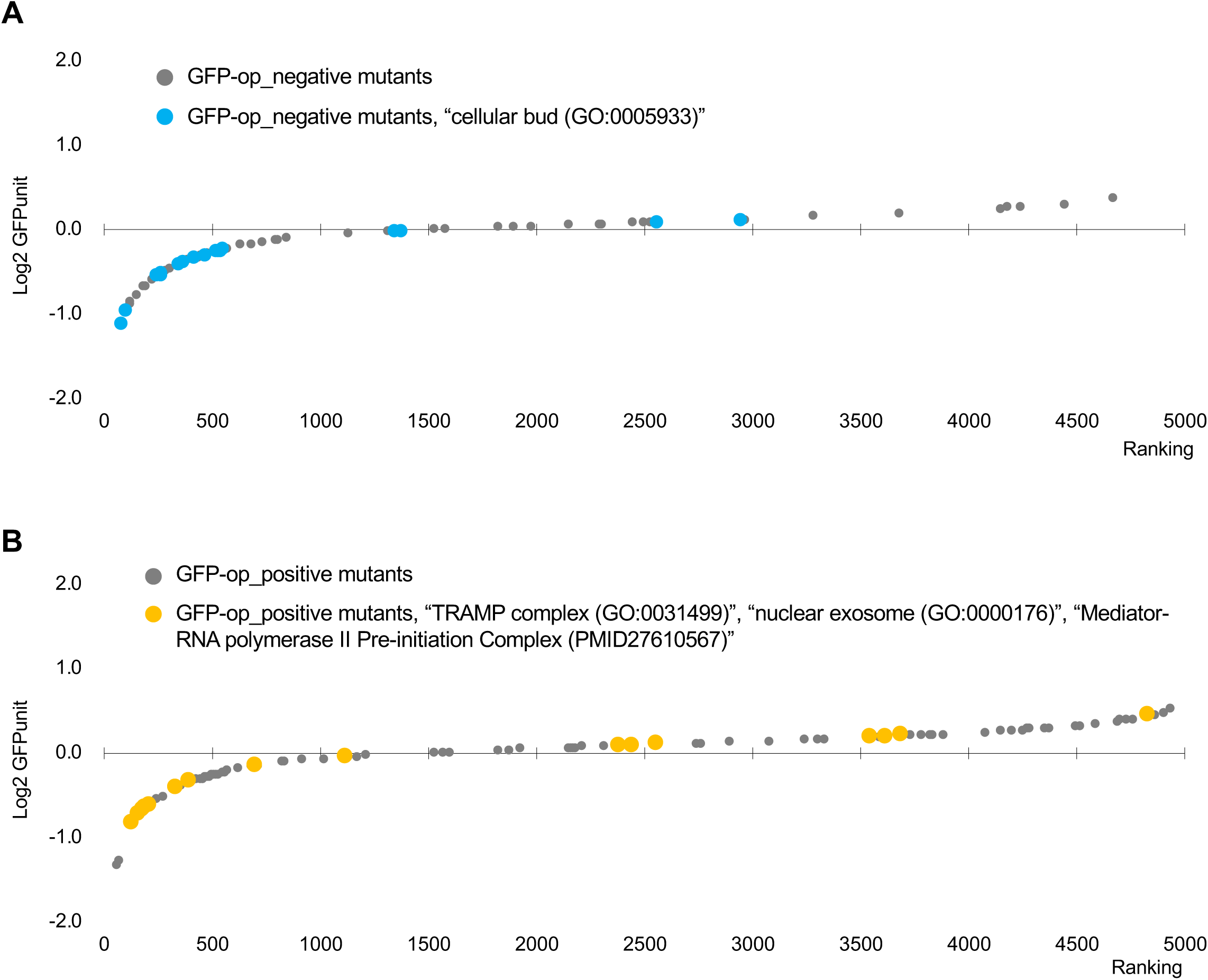
Distribution of GFPunits of mutants in specific GOs among mutants with GI with GFP-op. **A.** GFPunits of mutants in annotated to the GO term “cellular bud (GO:0005933)” among GFP-op_negative mutants. **B.** GFPunits of mutants annotated to the GO terms “TRAMP complex (GO:0031499),” “nuclear exosome (GO:0000176),” and “mediator-RNA polymerase II preinitiation complex (PMID27610567)” among GFP-op_positive mutants.

**Figure 5-S1.**
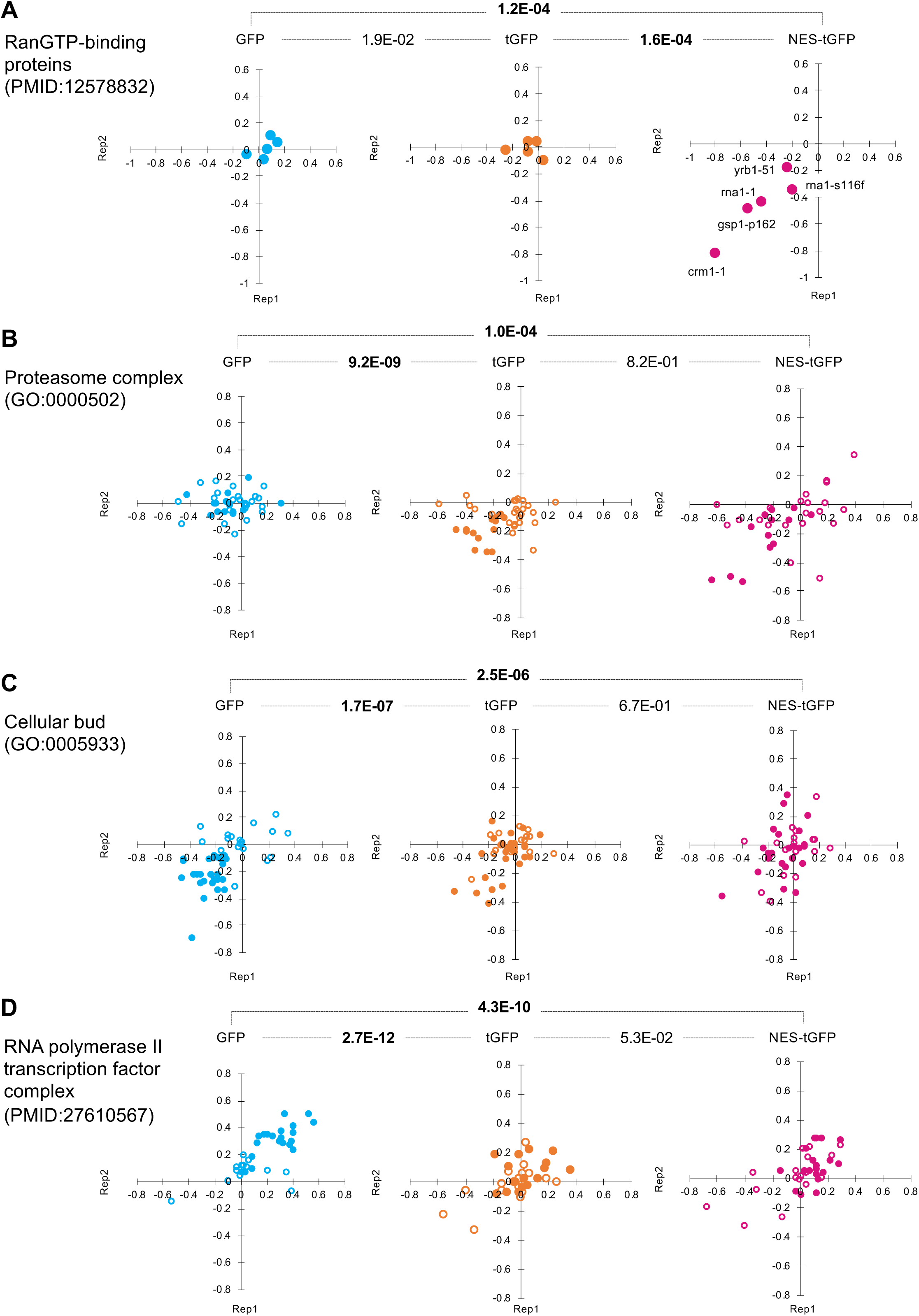
Distributions of GI scores of mutants in specific publications and GO. GI scores of mutants in the indicated publications and GO terms from the duplicated experiments on GFP-op, tGFP-op, and NES-tGFP are shown as scatter plots. The *p*-values of pair-wise t-test between experiments are shown. Bold letters indicate significant *p*-values (*p* < 0.001).

## Materials and Methods

### Strains and plasmids used in this study

The vector plasmid (pTOW40836), GFP-op plasmid (pTOW40836-TDH3_pro_-GFP), tGFP-op plasmid (pTOW40836-PYK1_pro_-NES-tGFP), and NES-tGFP-op plasmid (pTOW40836-PYK1_pro_-NES-tGFP) have been described previously (Kintaka *et al.*, 2016; Eguchi *et al.*, 2018). The deletion mutant collection and TS mutant collection have been described previously (Costanzo *et al.*, 2016). Yeast culture and transformation were performed as previously described (Amberg *et al.*, 2005). A synthetic complete (SC) medium without uracil (Ura) or leucine (Leu) was used for yeast culture.

### Query strains

Y7092 (MATa can1Δ::STE2pr-his5 lyp1Δ ura3Δ0 leu2Δ0 his3Δ1 met15Δ0) was used for the query train in the SGA. Y7092-E2-Crimson (MATa can1Δ::TDH3pr-E2-Crimson STE2pr-his5 lyp1Δ ura3Δ0 leu2Δ0 his3Δ1 met15Δ0) was used for the query strain in the SGA with the GFP fluorescent measurement experiment.

### Synthetic genetic array (SGA) and colony size analysis

SGA and colony size analysis were performed as previously described (Baryshnikova *et al.*, 2010). Briefly, an empty plasmid, and plasmids for overproducing GFP, tGFP, NES-tGFP were introduced into the deletion and TS mutant collections using robots to manipulate libraries in 1536-colony high-density formats. A query strain harboring each of the overexpression plasmids and each of the MATa mutant strains harboring a different genetic alteration were mated on YPD. Diploid cells were selected on plates containing both selection markers (YPD + G418 + clonNAT) found in the haploid parent strains. Sporulation was than induced by transferring cells to nitrogen starvation plates. Haploid cells containing all desired mutations were selected for by transferring cells to plates containing all selection markers (SC –His/Arg/Lys + canavanine + thialysine + G418 + cnonNAT) to select against remaining diploids. To analyze the growth of each deletion strain with the plasmids, all custom libraries were replicated to SC–LU plates and grown for three days at 30°C. The colony size was quantified and normalized. Then the genetic interaction (GI) scores were calculated using the formula. GI score (ε) = W_AB_ – W_A_ × W_B_, where W_AB_ is localized GFP plasmid/mutant fitness, W_A_ is localized GFP plasmid/WT fitness, and W_B_ is empty plasmid/mutant fitness. The GI scores were filtered using the defined confidence threshold (GI score, |ε| > 0.08, *p*-value that reflects both the local variability of replicate colonies (four colonies/ strain) and the variability of the strain sharing the same query or array mutation (*p* < 0.05). This filtered data set was used for all analyses. Initial GFP-op_positive 146 genes (147 mutants) contained genes involved in the His and Lys synthetic pathways. His and Lys (Arg) are used as marker genes for the SGA, and deletion mutants of *HIS, LYS*, and *ARG* genes should not grow in the SGA analysis. In fact, the colony sizes of these mutants in the vector control experiment were very small and were considered to be the carryover. We thus further isolated positively-interacting mutants by setting a threshold on the colony size of greater than 0.39 in the vector control experiment, selected according to the largest colony size (*ARG1*) among the *HIS, LYS*, and *ARG* mutants, to avoid the identification of false-positive GIs.

### Liquid growth measurement

Cellular growth was measured by monitoring OD595 every 30 min using a model 680 microplate reader (BioRad). The maximum growth rate was calculated as described previously (Moriya *et al.*, 2006). Average values and standard deviations were calculated from biological triplicates.

### GFP fluorescent measurement by Typhoon

Two colonies/strain from the SGA were picked up and replicated to SC–U plates, and grown for two days at 30°C. To detect the fluorescence of the colony, plates were scanned by laser (GFP at 488 nm and E2-Crimson at 532 nm) using Typhoon 9210 (Amersham Biosciences). The image data were analyzed using GenePix Pro Software (Molecular Devices). Each colony was segmented by a circle with the same diameter, the fluorescence per pixel was detected, and the medians of the fluorescence in the circle were calculated. To normalize the intensity by plate, all medians were divided by the plate average median for GFP and E2-Crimson. The ratios of GFP/RFP were calculated, and the averages of the two colonies were used.

### Clustering analysis

The GI scores of GFP, tGFP, and NES-tGFP were clustered into 15 clusters by the hierarchical clustering (average) method using R (https://www.r-project.org).

### Enrichment analysis

Enrichment analysis was performed using the gene list tool on the *Saccharomyces* genome database (yeastmine.yeastgenome.org/yeastmine/bag.do).

### Microscope observation

Log-phase cells were cultivated in SC–Ura medium. Cell images were obtained and processed using a DMI6000 B microscope and Leica Application Suite X software (Leica Microsystems). GFP fluorescence was observed using the GFP filter cube. Cellular DNA was stained with 100 μg/ml Hoechst 33342 (H3570, ThermoFisher) for 5 min and observed using the A filter cube.

### Quantification of GFP expression level

The total protein was extracted from log-phase cells with NuPAGE LDS sample buffer (ThermoFisher NP0007) after 0.2 N NaOH treatment for 5 min (Kushnirov, 2000). For each analysis, the total protein extracted from 0.1 optical density unit of cells OD600 1.0 was used. The extracted protein was labeled with Ezlabel FluoroNeo (WSE-7010, ATTO), as described in the manufacturer’s protocol, and separated by 4%–12% SDS-PAGE. Proteins were detected and measured using a LAS-4000 image analyzer (GE Healthcare) in SYBR–green fluorescence detection mode, and Image Quant TL software (GE Healthcare). The intensity of the 45kDa band corresponding to Pgk1 and Eno1/2 was used as the loading control. To detect GFP, the SDS-PAGE-separated proteins were transferred to a PVDF membrane (ThermoFisher). GFP was detected using an anti-GFP antibody (11814460001, Roche), a peroxidase-conjugated second antibody (414151F, Nichirei Biosciences), and a chemiluminescent reagent (34095, ThermoFisher). The chemiluminescent image was acquired with a LAS-4000 image analyzer in chemiluminescence detection mode (GE Healthcare). For the estimation of relative GFP levels, the intensities of corresponding GFP bands were normalized using the loading control described above.

### High-dimensional morphological analysis

Morphological data of cells cultured were acquired as previously described (Ohya *et al.*, 2005). Briefly, logarithmic-phase BY4741 cells harboring plasmids grown in SC–Ura medium were fixed and were triply stained with FITC-ConA, rhodamine-phalloidin, and 4,6-diamidino-2-phenylindole to obtain fluorescent images of the cell-surface mannoprotein, actin cytoskeleton, and nuclear DNA, respectively. Images of at least 200 individual cells were acquired and processed using CalMorph (version 1.2). All of the statistical analyses were performed with R. To statistically test the morphological differences among four strains, we conducted one-way ANOVA of the generalized linear model (GLM) for each of 501 morphological parameters. Probability density functions (PDFs) and accompanying link functions in the GLM were assigned to each trait as described previously (Yang *et al.*, 2014). Difference of the four strains (n = 5) was incorporated as the explanatory variable into the linear model. We assessed a dispersion model among the strains in the linear models for the 501 parameters by Akaike information criterion (AIC) and set 110 models (parameters) as a different dispersion model because of lower AIC than that of a single dispersion model. Applying one-way ANOVA among the four strains to all 501 parameters, 51 of the 501 parameters were found to differ significantly at false discovery rate (FDR) = 0.01 by the likelihood ratio test (**Likelihood ratio test in Data S7**). Maximum likelihood estimation, likelihood ratio test, and the estimation of FDR were performed using the gamlss, lrtest, and qvalue functions in the gamlss (Stasinopoulos and Rigby, 2007), lmtest (Zeileis and Hothorn, 2002), and qvalue (Storey, 2002) R package. By Wald test at FDR = 0.01, 16, 17, and 24 of the 501 traits were detected to have a significant difference from wild-type in GFPy66g, Gpm1, and Gpm1-m, respectively (**Q value of Wald test in Data S7**). Of the 16 parameters detected in GFPy66g, 14 parameters were grouped into four independent morphological features by four principal components (explaining 60% of the variance) extracted from principal component analysis for the Z values of 109 replicates of *his3Δ* (Suzuki *et al.*, 2018) as described previously (Ohnuki *et al.*, 2012), and were used for the illustration of morphological features (**Figure 6F, Morphological features in in Data S7**).

## Acknowledgements

KAKENHI 17H03618, KAKENHI 15KK0258 We thank members of the Moriya lab, the Boone lab and the Andrews lab for advice and helpful discussions.

## Supplementary Datasets and Tables

- **Data S1 (Data_S1_GFP_SGA_raw_data.xlsx)** Raw data of GFP-op SGA analysis; associated with Figure 2A-C, 2-S1A-D, 2-S5A, and 3A-C.
- **Data S2 (Data_S2_GFPunit.xlsx)** Raw data of GFP expression analysis under GFP-op; associated with Figure 4B, 4C, and 4-S1A, and 4-S1B.
- **Data S3 (Data_S3_NES-tGFP_SGA_raw_data.xlsx)** Raw data of NES-tGFP-op SGA analysis; associated with Figure 5-S1A-D.
- **Data S4 (Data_S4_tGFP_SGA_raw_data.xlsx)** Raw data of tGFP-op SGA analysis; associated with Figure 5-S1A-D.
- **Data S5 (Data_S5_GFP_isolated_mutants.xlsx)** Isolated GFP-op_positive and _negative mutants by this study; associated with Figure 2D, 4B, and 4C.
- **Data S6 (Data_S6_GFP_tGFP_NES-tGFP_isolated_cluster.xlsx)** Isolated mutants with GFP, tGFP, and NES-tGFP SGA analysis, and the result of clustering analysis; associated with Figure 5C and 5D.
- **Data S7 (Data_S7_Morphological_Phenotyping.xlsx)** Whole dataset of morphological phenotyping of overexpressing strains; associated with Figure 6A-D.
- **Table S1 (Table_S1_Enrichement SGA_GFP.xlsx)** Enrichment analysis of genes isolated in GFPop SGA analysis; associated with Figure 2D.
- **Table S2 (Table_S2_Enrichiment_GFPunit.xlsx)** Enrichment analysis of genes isolated by GFP expression level; associated with Figure 4B, 4C, 4-S1A, and 4-S1B.
- **Table S3 (Table_S3_Clusters_Genes_Enrichement.xlsx)** Enrichment analysis of genes in each cluster in Figure 5C and 5D.

